# Parietal Cortex Integrates Object Orientation and Saccade Signals to Update Grasp Plans

**DOI:** 10.1101/758532

**Authors:** Bianca R. Baltaretu, Simona Monaco, Jena Velji-Ibrahim, Gaelle N. Luabeya, J. Douglas Crawford

## Abstract

Coordinated reach-to-grasp movements are often accompanied by rapid eye movements (saccades) that displace the desired object image relative to the retina. Parietal cortex compensates for this by updating reach goals relative to current gaze direction, but its role in the integration of oculomotor and visual orientation signals for updating *grasp* plans is unknown. Based on a recent perceptual experiment, we hypothesized that inferior parietal cortex (specifically supramarginal gyrus; SMG) integrates saccade and visual signals to update grasp plans in more superior parietal areas. To test this hypothesis, we employed a functional magnetic resonance adaptation paradigm, where saccades sometimes interrupted grasp preparation toward a briefly presented object that later reappeared (with the same/different orientation) just before movement. Right SMG and several parietal grasp areas, namely left anterior intraparietal sulcus (aIPS) and bilateral superior parietal lobe (SPL), met our criteria for transsaccadic orientation integration: during movement preparation, they showed task-dependent saccade modulations and, during grasp execution, they were specifically sensitive to changes in object orientation that followed saccades. Finally, SMG showed enhanced functional connectivity with both prefrontal saccade areas (consistent with oculomotor input) and aIPS / SPL (consistent with sensorimotor output). These results support the general role of parietal cortex for the integration of visuospatial perturbations, and provide specific cortical modules for the integration of oculomotor and visual signals for grasp updating.

**Significance Statement:** The cortical mechanisms that update reach goals during eye movements are well documented, but it is not known how object features are linked to oculomotor signals when updating grasp plans. Here, we employed functional magnetic resonance imaging adaptation (fMRIa) and functional connectivity analysis to identify a cluster of inferior parietal (supramarginal gyrus) and superior parietal (intraparietal and superior parietal) regions that show functional connectivity with frontal cortex saccade centers, and also show saccade-specific modulations during unexpected changes in object / grasp orientation. This provides a network - complementary to the goal updater network - that integrates visuospatial updating into grasp plans, and may help explain some of the more complex symptoms associated with parietal damage such as constructional ataxia.

## Introduction

We inhabit a dynamic visual environment, where brain and behavior must constantly compensate for changes in relative visual location induced by our own motion and/or external changes. For example, parietal cortex is thought to play an important role in updating reach goals in response to both unexpected changes in object location (1) and internally driven changes in eye position (2–4). The latter often compensates for rapid eye movements (saccades), allowing reaches toward targets that are no longer visible (4, 5), and more precise aiming to visible targets (6, 7). However, successful object interaction often requires more than transporting the hand towards the target, it also requires grasping: shaping of the hand to fit specific object attributes, such as shape and orientation (8–10). Reach transport and hand configuration must be intimately coordinated through space and time for successful grasp ((11); see (12) for review). Likewise, intended grasp location and orientation must remain linked and updated during saccades (13, 14). However, to date, the cortical mechanisms that integrate saccade and object features for grasp updating have not been studied.

Clues to visual feature updating for grasp might be gleaned from studies of transsaccadic perception: the comparison and integration of visual information obtained before and after a saccade (15–17). Transcranial magnetic stimulation studies suggest that the frontal eye field (FEF) provides the saccade efference copy for transsaccadic integration of stimulus orientation, and that posterior parietal cortex is also involved (18, 19). Recently, a functional magnetic resonance imaging adaptation (fMRIa) paradigm showed that the inferior parietal lobe (specifically, the supramarginal gyrus; SMG) is specifically sensitive to transsaccadic changes in visual stimulus orientation (20). Human SMG probably expands functionally and anatomically into the lateral intraparietal cortex in the monkey, which contains a mixture of saccade, visual feature, and spatial updating signals (21–23). Since the inferior parietal cortex is thought to play an intermediate role in perception and action (24), spanning both ventral and dorsal stream visual functions (25), we hypothesized that SMG might also play a role in updating stimulus orientation for grasp planning across saccades.

Feature updating, however, influences behavior only when it results in the updating of sensorimotor plans. Several parietal areas have been implicated in the visuomotor transformations for grasp, including the anterior intraparietal sulcus (aIPS) (21, 26), superior parietal cortex (SPL) (22, 27), and superior parieto-occipital cortex (SPOC) (28, 29). Transcranial magnetic stimulation experiments suggest that SPOC is involved in early visuomotor transformations for setting reach goals (30, 31), whereas more anterior areas along the intraparietal sulcus have also been implicated in updating grasp plans in response to external perturbations in object shape and orientation (32, 33). However, it is not known if these (or other) areas are involved in the integration of saccade and orientation signals for transsaccadic updating when planning a grasping movement.

Based on this background information, we hypothesized that SMG and areas in the parietal grasp network provide the visuomotor coupling for transsaccadic grasp updating, by integrating visual feature input with internal saccade signals, originating from frontal cortex. To test this model, we performed a series of experiments in an MRI suite equipped with an eye tracker and a rotatable grasp stimulus that could be presented in complete darkness (Fig. 1*A*). Specifically, we merged two previously employed event-related fMRIa paradigms for transsaccadic integration (20) and grasp planning (34), respectively (Fig. 1*B*). We applied a set of criteria to identify areas involved in the integration of eye position and visual orientation changes for grasp updating: 1) these areas should be specifically sensitive to transsaccadic changes in visual orientation during grasping movements (20) (Fig. 1 *C1*), 2) they should show saccade modulations during grasp preparation (Fig. 1 *C2*), and 3) these modulations should be task-specific, especially in areas associated with the sensorimotor control of grasp (Fig. 1 *C3*). Finally, during grasp updating, these areas should show stronger functional connectivity for saccades than fixation, both with each other, and with the putative source of an oculomotor signal originating in the cortical saccade generator.

**Figure 1.**
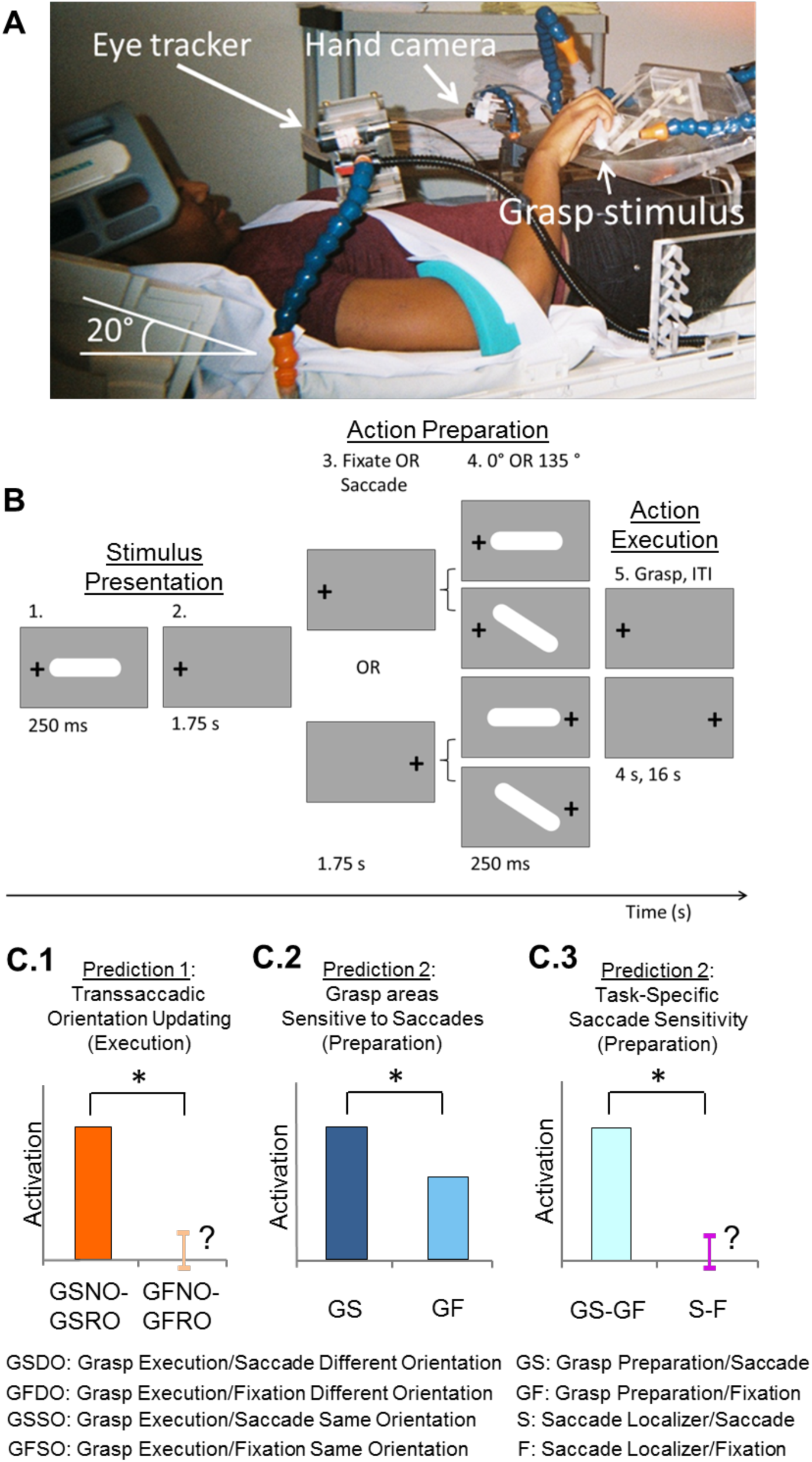
Experimental set-up, paradigm, and predictions. ***A*.** Set-up of the experiment, showing participant lying supine on MRI table with head tilted at 20° under the head coil, along with MRI-compatible eye tracker for right eye and hand tracker. Participants rested their hand on the abdomen in a comfortable position and were asked to transport their hand to the platform to grasp an oriented 3D bar only when required to do so; a strap across the torso was used to ensure minimal-to-no movement of the shoulder and arm during transportation of the hand to the platform. The blue stalk above the platform was used to illuminate the central grasp object, whereas those to the left and right contained LEDs and were used to ensure fixation of gaze. ***B*.** Stimuli and task. An example of an initial trial condition is shown (0° grasp bar, gaze left) followed by the four possible conditions that might result: Fixate / Different Feature, Fixate / Same Feature, Saccade / Different Feature; and Saccade / Same Feature). Each trial lasted 24 seconds and was comprised of three major phases: 1) *Stimulus Presentation*, during which the grasp object was illuminated in one of two possible orientations (0° or 135°) and gaze could be left or right; 2) *Action Preparation*, when participants maintained fixation on the same LED as in the previous phase (Fixate condition) or they made a saccade to the opposite LED (Saccade condition) – the object was illuminated a second time at the end of this phase and was presented either in the Same orientation as in phase 1 (0° if the initial was 0° or 135° if the initial orientation was 135°; Same condition) or at a Different orientation (0° if the initial was 135° or vice versa; Different condition); and 3) *Action Execution*, which required participants to grasp the oriented object within 4 s and then, return to rest (only the first 2 s were used for analysis). This was followed by an inter-trial interval of 16 s. ***C*.** The possible predictions for sensitivity to saccade signals in grasp areas in three conditions. *C.1*. The first prediction suggests that, during the *Action Execution* phase, cortical regions that specifically update object orientation across saccades should show a greater difference in activity between the Same and Different orientation conditions in the Grasp Saccade condition, as compared with the Same – Different orientation difference in the Grasp Fixate condition (GSDO, GSSO, GFDO, GFSO, respectively). *C.2*. The second prediction indicates that, if a grasp area is modulated by saccade signals, the BOLD activity should be greater for the Saccade condition (Grasp Saccade condition, GS), as compared with the Fixate condition (Grasp Fixation condition, GF). *C3*. The third prediction tests whether modulations due to saccade signals during the grasp *Action Preparation* phase (*C.2*) are specific to grasp-related activity. This predicts a greater difference between the Saccade and Fixate conditions in the grasp experiment compared to a separate saccade localizer that only required participants to either saccade between our two LEDs or fixate on one of the LEDs ((Grasp Saccade - Grasp Fixate) > (Saccade – Fixate); GS - GF > S - F).

## Results

### Grasp Planning and Saccade Modulations during Action Preparation

As indicated in Fig. 1*B*, within each trial, there were three key phases: *Stimulus Presentation* (which begins with the original grasp stimulus orientation), *Action Preparation* (which included a saccade in 50% of trials, and ends with a Novel or Repeat stimulus orientation that also acts as a ‘go’ signal), and *Action Execution* (where the actual reach and grasp occurs). By design, we expected brain activation to be dominated by: 1) visual signals during the *Stimulus Presentation* phase, 2) grasp preparation, saccade, and spatial updating signals during the *Action Preparation* phase, and 3) grasp motor signals and (in the case of Novel stimuli) grasp orientation updating during the *Action Execution* phase of this task. We begin with an overview of the activation in the *Action Preparation* phase, where one might expect to find events related to the saccade-related updating of the original grasp stimulus.

Various studies have shown that humans can remember stimulus properties for several seconds, and use these to plan action until a ‘go’ signal is provided (35, 36). To isolate activity related to grasp preparation, we contrasted activity from the *Action Preparation Phase* (between 1^st^ and 2^nd^ stimulus; Fig. 1*B*) of Grasp Fixation trials against baseline activity (Fig. 2*A*). This revealed activation in a parietofrontal network, including right SMG and several well-established reach/grasp areas: aIPS, lateral SPL (lSPL), precentral gyrus (PCG; corresponding to primary motor cortex), and dorsal / ventral precentral sulcus (PCSd/ PCSv; likely portions of these areas corresponding to dorsal and ventral premotor cortex, respectively) (12, 22, 37). In short, the initial stimulus evoked massive preparatory activity in the grasp network.

**Figure 2.**
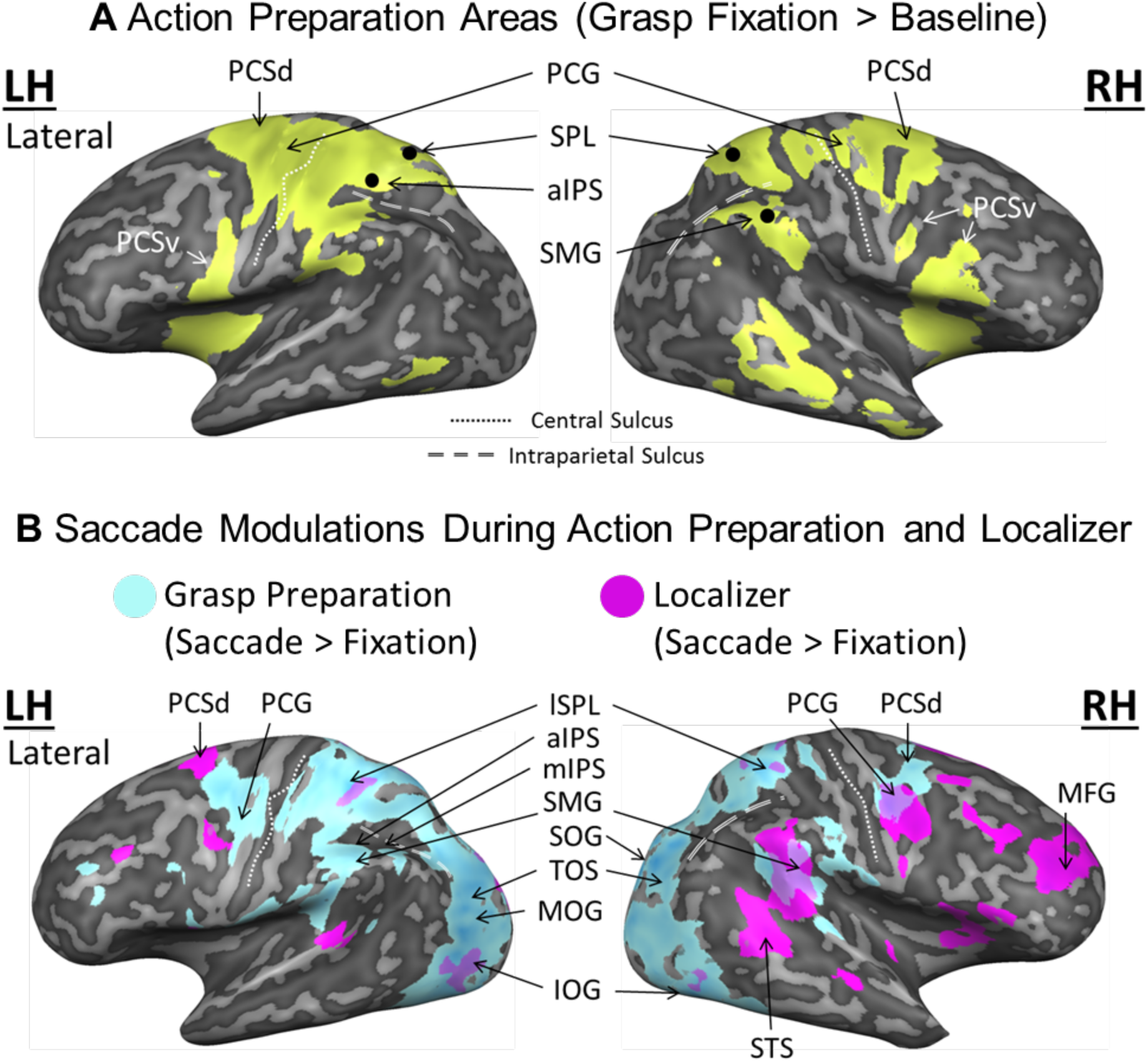
Lateral view of (A) reach/grasp cortical regions and (B) saccade modulations during *Action Preparation. **A***. Shown is an inflated brain rendering of an example participant (left and right hemispheres from the lateral view, respectively). An activation map obtained using an RFX GLM (n=13) is shown for the contrast, Grasp Fixation > Baseline (chartreuse). *Abbreviations*: PCSd: dorsal precentral sulcus, PCSv: ventral precentral sulcus, PCG: precentral gyrus, aIPS: anterior intraparietal sulcus, SPL: superior parietal lobe, SMG: supramarginal gyrus. ***B*.** Activation maps for a Saccade > Fixate contrast obtained using an RFX GLM (n=13) on grasp experiment data (sky blue) and on a separate saccade localizer (fuchsia) were overlaid onto an inflated brain rendering from an example participant (left and right hemispheres shown in the lateral views). *Abbreviations:* PCSd: dorsal precentral sulcus, PCSv: ventral precentral sulcus, PCG: precentral gyrus, lSPL: lateral superior parietal lobe, aIPS: anterior intraparietal sulcus, mIPS: middle intraparietal sulcus, SMG: supramarginal gyrus, SOG: superior occipital gyrus, TOS: transverse occipital sulcus, MOG: middle occipital gyrus, IOG: inferior occipital gyrus, STS: superior temporal sulcus.

To detect if grasp planning was also modulated by saccades, we compared grasp saccade trials to grasp fixation trials during the *Action Preparation* phase (Fig. 2*B,* sky blue areas), and compared this to activity from our saccade localizer task (Fig. 2*B*, fuchsia areas). These two contrasts produced overlap in some cortical regions (e.g., right frontal cortex and SMG), but saccades also produced extensive superior parietal and occipital modulations in the grasp task, including traditional grasp areas like aIPS and SPL. However, these additional modulations could be related to various functions, such as updating reach *goals* (2, 3), general aspects of eye-hand coordination (38), or expected sensory feedback (39). To identify activity specific to *transsaccadic grasp updating*, we used our *a priori* predictions (Fig. 1 *C.1, 2, 3),* as shown in the following analyses.

### Prediction 1: Saccade-Specific Sensitivity to Stimulus Orientation Changes

The first step was to determine regions that are specifically involved in transsaccadic grasp updating. If our participants incorporated original object orientation into short term memory and uses this for grasp planning (34, 35), update this information across saccades (17, 20) and then update this again when they saw the final object orientation (40, 41), the cortical response to the second stimulus should be modulated by the orientation of the first stimulus (34), and some of these modulations should depend on changes in eye position. Specifically, we predicted that these areas should show an increased response to orientation changes in the Grasp/Saccade condition and little or no increase in the Grasp/Fixation condition (Figure 1 C1). Alternatively, if participants ignored the initial stimulus and waited for the final stimulus to plan the grasp, these modulations should not occur.

Based on previous literature, we hypothesized that this might involve both right SMG (20) and the intra/superior parietal grasp network (32, 33). To test this, we applied prediction 1 (Fig. 1 *C.1*) to the *Action Execution* interval following stimulus re-presentation, looking for visual/motor areas only sensitive to changes in grasp orientation (Novel versus Repeat orientation) that follow saccades. Specifically, we used a voxelwise contrast applied to the trials wherein a saccade or fixation occurred (i.e., (Grasp Saccade Novel Orientation > Grasp Saccade Repeat Orientation) > (Grasp Fixation Novel Orientation > Grasp Fixation Repeat Orientation)).

As shown in Fig. 3*A*, this contrast predominantly identified several parietal areas that where responses to the second stimulus were modulated by initial stimulus orientation in a saccade-specific fashion. In particular, right SMG, left aIPS, and bilateral SPL showed saccade-specific responses to changes in stimulus orientation. All four of these areas passed cluster threshold correction (see Methods for details and Table 1 for Talairach coordinates). Fig. 3*B* shows the same result, but presented in a more quantitative format (β-weights extracted from voxels of peak activation) designed to enable a direct visual comparison with prediction 1 (Fig. 1 *C.1)*. We will henceforth refer to these four sites as putative grasp updating sites.

**Figure 3.**
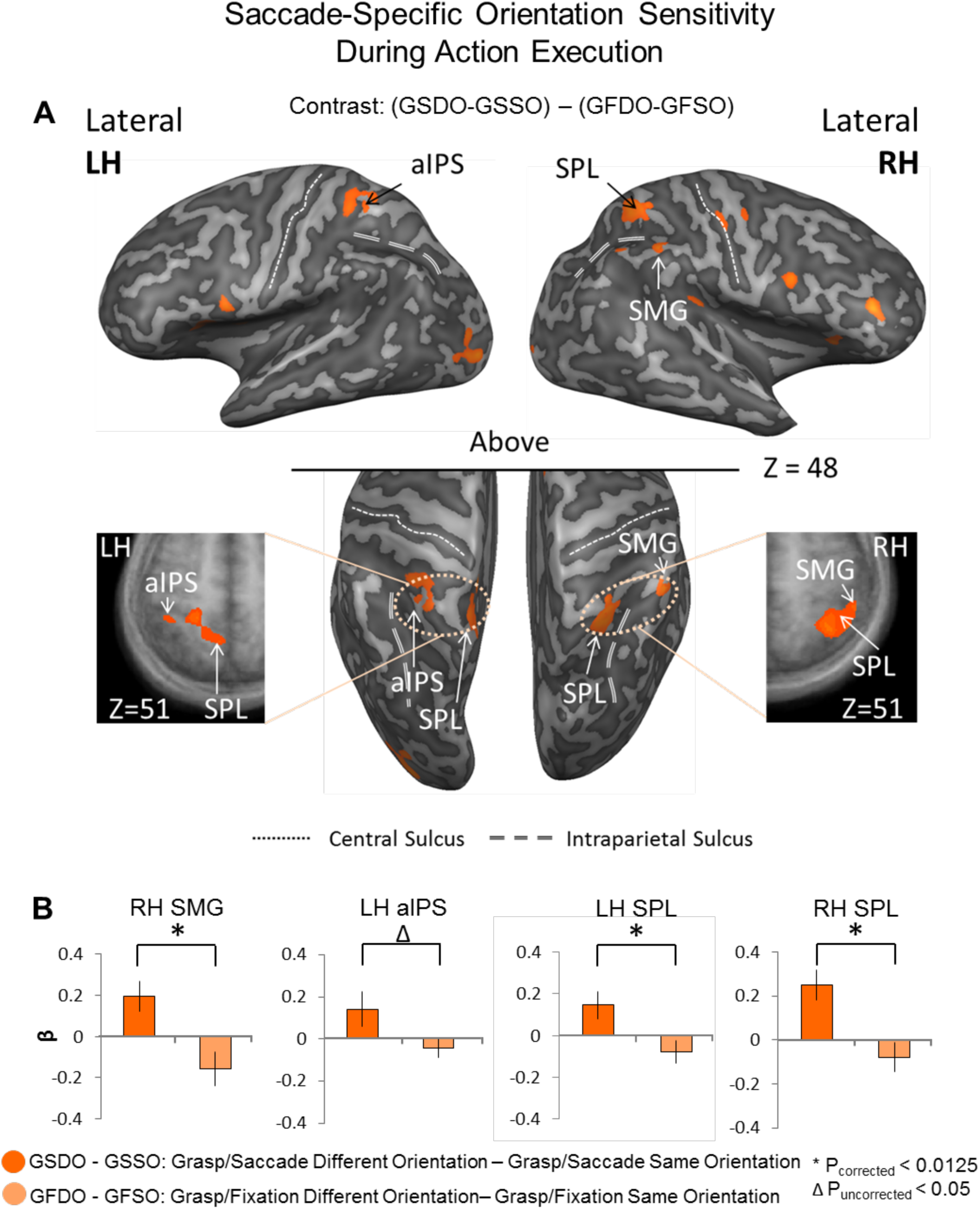
Post-saccadic Different vs. Same orientation responses during *Action Execution*. ***A.*** Voxelwise statistical map overlaid onto inflated brain rendering of an example participant obtained using an RFX GLM (n=13) for Different > Same in the Grasp Saccade condition as compared with the Grasp Fixate condition (p<0.05). Top panels show the lateral views of the inflated brain rendering on which can be seen activation in right superior lateral lobe (SPL) and supramarginal gyrus (SMG). In the middle, bottom panels, the top view of the left and right hemispheres can be seen, which display activation also in the left anterior intraparietal sulcus (aIPS) and SPL. The left and rightmost panels contain transverse slices through the average brain of all the participants onto which the activation in these five regions can be viewed in more detail. These results (that the final motor plan was modulated by the initial stimulus orientation) contradict the notion that participants waited for the second stimulus orientation to begin action planning. Instead, they show that an orientation-specific action plan was formed immediately, and then updated when the second stimulus was presented. ***B.*** Bar graphs of β-weights plotted for the difference between the Grasp/Saccade Different and Same orientation conditions (dark orange) versus the difference between the Grasp/Fixation Different and Same conditions (light orange). The small, variable Grasp/Fixation results are analogous to the results of sensory adaptation studies, where both repetition suppression and enhancement effects have been observed (59). Data were extracted from peak voxels from the transsaccadic regions shown in A. Statistical tests were carried out on β-weights extracted from peak voxels on these areas in order to test Prediction 1. Values are mean ± SEM analyzed by dependent *t* test. * indicates a statistically significant difference between the GS and GF β-weights during the *Action Preparation* phase (Bonferroni corrected at a p<0.0125). Δ indicates an uncorrected significant difference between the GS and GF β-weights during the preparatory period (not Bonferroni corrected, p<0.05).

**Table 1.** Talaraich coordinates for regions-of-interest extracted from the *Action Execution* phase.

| ROI Name | Talairach coordinates |  |  |  |  |  | ROI size |
| --- | --- | --- | --- | --- | --- | --- | --- |
|  | x | y | z | Std x | Std y | Std z | *n* voxels |
| **LH aIPS** | -38 | -41 | 53 | 2.3 | 1.6 | 1.9 | 217 |
| **LH SPL** | -13 | -51 | 54 | 2.4 | 2.3 | 2.7 | 642 |
| **RH SMG** | 49 | -40 | 48 | 2.0 | 2.2 | 2.1 | 361 |
| **RH SPL** | 35 | -49 | 53 | 2.8 | 2.8 | 2.7 | 875 |

### Predictions 2 and 3: Site-Specific Saccade Modulations and Task Specificity

To directly examine the influence of saccades on the four cortical sites identified in the previous section, we examined *Action Preparation* phase activity of our task (prediction 2) and compared this to our separate saccade localizer data (prediction 3). Panels *A* and *B* of Figure 4 show the locations of the peak voxels from our putative grasp updating sites, superimposed on the overall preparatory activity during fixation only, and saccade modulations in our task and saccade localizer, respectively (derived as in Fig. 2). All four sites (right SMG, left aIPS, and bilateral SPL) fell within regions of grasp preparation (Fig. 4*A*), as well as within, or bordering on, regions of saccade modulation (Fig. 4*B*). To test saccade sensitivity in these regions (during *Action Preparation*), we applied prediction 2 on β-weights extracted from these locations (Fig. 4*C*). All four regions showed significantly higher preparatory activity in the presence of saccades, although SMG did not survive correction for multiple comparisons. To test the task specificity of these modulations, we applied prediction 3, i.e., we tested if these locations showed saccade modulations during grasp preparation, but not during saccades alone (Fig. 4*D*). In this case, only aIPS and bilateral SPL showed significant task specificity. This suggests a progression of task-specificity from SMG to the more motor areas.

**Figure 4.**
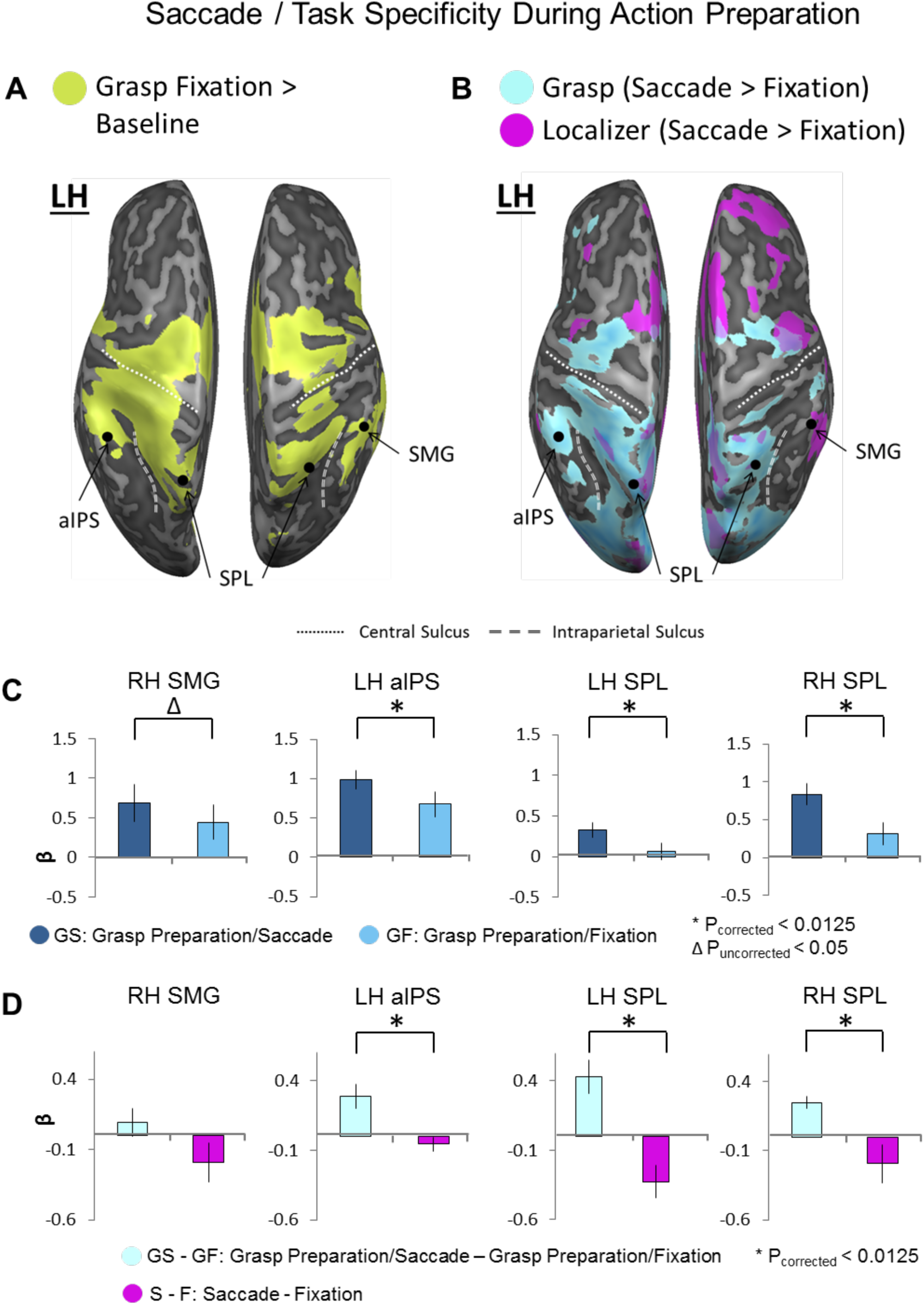
Location of putative transsaccadic reach updating sites (from Fig. 3) relative to grasp areas (A) and saccade modulations (B) during *Action Preparation*, followed by prediction tests 2 (C) and 3 (D). ***A*.** Shown is an inflated brain rendering of an example participant (left and right hemispheres viewed from above, respectively). An activation map obtained using an RFX GLM (n=13) is shown for the contrast, Grasp Fixation > Baseline (chartreuse). The four transsaccadic regions from the *Action Execution* phase are overlaid onto this *Action Preparation* activation. aIPS: anterior intraparietal sulcus, SPL: superior parietal lobe, SMG: supramarginal gyrus. ***B*.** Activation maps for a Saccade > Fixate contrast obtained using an RFX GLM (n=13) on grasp experiment data (sky blue) and on a separate saccade localizer (fuchsia) were overlaid onto an inflated brain rendering from an example participant (left and right hemispheres shown from a bird’s eye view). These overlaid activation maps allow for comparison of which cortical regions respond to saccade signals in a grasp task-specific manner. *Abbreviations:* aIPS: anterior intraparietal sulcus, SPL: superior parietal lobe, SMG: supramarginal gyrus. ***C.*** Bar graphs of β-weights plotted for Grasp Saccade conditions (dark blue) versus Grasp Fixation conditions (light blue) from all thirteen participants. Data were extracted from peak voxels from the transsaccadic regions represented by the black dots above in *A* and *B* in order to test prediction 2. Values are mean ± SEM analyzed by dependent *t* test. ***D.*** Bar graphs of β-weights plotted for Grasp Saccade conditions (pale blue) versus Grasp Fixation conditions (magenta). Data were extracted from peak voxels from the transsaccadic regions shown in Fig. 2A and *B,* which compared for only the ten participants whose data were analyzed for the saccade localizer. Statistical tests were carried out on β-weights extracted from peak voxels in these areas in order to test prediction 3. Values are mean ± SEM analyzed by dependent *t* test. * indicates a statistically significant difference between the GS and GF β-weights during the preparatory period (Bonferroni corrected at a p<0.0125). Δindicates an uncorrected significant difference between the GS and GF β-weights during the preparatory period (not Bonferroni corrected, p<0.05).

### Functional Connectivity of SMG with Saccade and Grasp Areas

Our analyses so far have confirmed our perceptual updating result for SMG (20), and extended this function to sensorimotor updating in aIPS and SPL for grasp; but, do these regions participate in a coherent functional network for grasp updating? Based on our previous finding that right SMG is active for perceptual orientation updating (20), and its re-appearance in the current grasp task, we hypothesized that SMG is a key hub for updating visual orientation across saccades, and that it would communicate with both saccade regions (for signal input) and grasp regions (for signal output) during our grasp task. To do this, we identified a seed region within the right SMG from our independent saccade localizer data, and performed a psychophysiological interaction (PPI) analysis to examine which areas showed increased functional connectivity for saccade as compared with fixation trials with SMG during *Action Preparation* (Fig. 5*A-C*). This resulted in three sites that survived cluster threshold correction: right PCSd (likely a portion corresponding to FEF), left medial, superior frontal gyrus (likely the supplementary eye field, SEF), and SPL (including a region that overlaps with aIPS).

**Figure 5.**
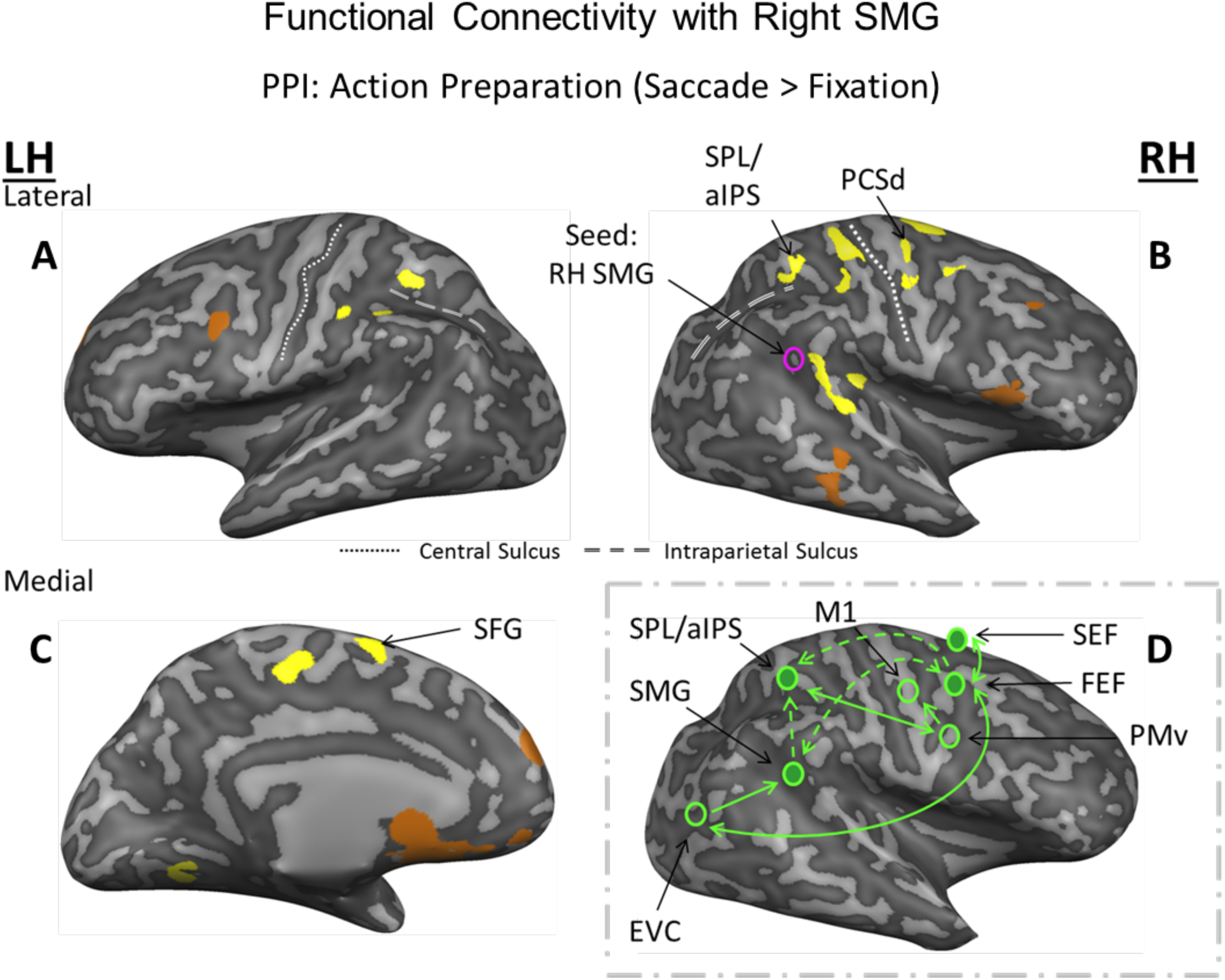
Functional connectivity network involved in processing saccade signals during *Action Preparation*. ***A-C.*** Using a Saccade > Fixation contrast and the right supramarginal gyrus (SMG) as a seed region, psychophysiological interaction is shown in the activation maps (yellow for positive correlation; copper for negative correlation) overlaid onto the inflated brain renderings of an example participant. Right frontal eye field (FEF), SPL (that extends into the anterior intraparietal sulcus, aIPS) and left supplementary eye field (SEF) show significant, cluster-corrected positive correlation with right SMG. Only areas that passed a p<0.05 and cluster threshold correction are labeled. ***D.*** A potential network for the communication between right SMG and other saccade and grasp regions.

## Discussion

In this study, we set out to identify the cortical areas associated with updating grasp plans during changes in gaze direction and/or object orientation. We reasoned that, in order to perform this function, the brain would have to integrate saccade signals in areas sensitive to visual orientation and/or grasp orientation updating. To identify these areas, we applied three specific criteria: transsaccadic sensitivity to orientation changes during grasp execution, sensitivity to intervening saccades during motor preparation, and task specificity in these modulations, at least in the more superior parietal grasp motor areas. We found four areas that met these criteria: right SMG, an area previously implicated in transsaccadic orientation perception (20), and three more dorsal areas that are associated with grasp correction (18, 30). Finally, with the use of task-related functional connectivity analysis with area SMG, we identified a putative network for saccades that includes parietal and prefrontal regions.

### Transsaccadic Updating of Object Orientation for Grasp

In a previous study, we found that the right anterior inferior parietal lobe (SMG) is involved in transsaccadic comparisons of object orientation for perception (20). Here, we hypothesized that SMG would contribute to feature updating for grasp execution, whereas some part of other areas involved planning/updating grasp orientation (26, 42, 43) would also be involved in the transsaccadic updating of orientation for grasp preparation. To test this, we compared orientation change specificity for saccades versus fixation during *Action Execution*, and found four areas (right SMG, left aIPS, and bilateral SPL) that fit this criterion and passed our standard statistical criteria. We further found that all of these areas were modulated by saccades during grasp preparation, although the motor task specificity of these modulations was clearer in aIPS and SPL. Finally, the laterality of these responses was consistent with our hypothesis, i.e., right SMG being consistent with the general role of right parietal cortex in spatial awareness (44), whereas left aIPS was opposite to the motor effector uses (the right hand). This supports a general-purpose role for right SMG in the transsaccadic updating of object orientation, and adds a more unique role for aIPS and SPL in updating grasp orientation.

SMG is an area that has largely been implicated in perception tasks, such as those requiring spatial processing of orientation (45) and visual search (46), or those requiring crossmodal spatial attention (47). In contrast, SPL has been implicated in both saccade- and grasp-related populations (27) that make it an ideal site to respond to changes in retinal visual information about the position and orientation of an object. This may alter any hand preshaping signals that will be sent from SPL (48) to PMd (49, 50), which possesses both mixed saccade-and-reach or reach-only populations of neurons (27). Finally, aIPS is sensitive to object orientation information for grasp (26, 28, 51, 52). aIPS appeared twice in our analysis: first in Fig. 3, near the coordinates provided in some previous studies (34, 48, 53) and second, clustered with SPL in our network analysis (Fig 5). It is thought that populations of neurons in aIPS may process object features such as its orientation in order to ultimately shape and orient the hand to match the object’s shape and orientation (42). Information related to grasping is then proposed to travel to PMv to engage specific reach/grasp-related neuronal populations to generate motor commands (27, 50, 54). Thus, our result appears to be consistent with the known functions of these areas, and extends our understanding of how these functions might be linked to update grasp signals in the presence of saccades.

### A Putative Network for Transsaccadic Updating of Grasp Plans

An important goal for this study was to understand how distributed cortical regions might work as a network to update grasp plans during saccades. Based on the computational requirements of this function, we hypothesized that such a network should involve: 1) areas specific to transsaccadic updating of orientation features, 2) saccade areas for oculomotor input, and 3) and grasp updating areas for motor output. Given our previous (20) and current results, we hypothesized that right SMG would play the first role (i.e., here, it would update object features across saccades during the *Action Preparation* phase so that these could be spatially integrated with new visual information for *Action Execution*), and chose this as the seed region for our functional connectivity analysis. As described in the Introduction, we expected prefrontal saccade areas to play the second role, and parietal grasp areas to provide the final role (based on our current results, aIPS/SPL). Indeed, this analysis revealed a functional network for saccades versus fixation involving right SMG, right SPL, right aIPS, right PCSd, and the left superior frontal gyrus. Taken together with the overlapping areas that fit the previous 3 criteria, this suggests a saccade-dependent network with the specific properties needed for updating grasp orientation.

PPI analysis does not provide directionality, but based on the functional requirements of the task and known physiology of these areas, we conceptualized this network as shown in Fig. 5*D*. PCSd likely corresponds to the right FEF (55, 56). The FEF is a key component of the cortical saccade generator (56), and is known to provide reentrant feedback to earlier visual areas (57, 58). The superior frontal gyrus likely corresponds to the supplementary eye field (56, 59), which has reciprocal connections with FEF. Thus, FEF/SEF could be the source of saccade signals for SMG and the entire network. As discussed above, aIPS (48) and SPL are implicated in grasp planning / corrections, show saccade signals (27, 60), and of course were already implicated in transsaccadic grasp updating in our other analyses. Thus, this putative network appears to possess all of the signals and characteristics that one would expect to find in a transsaccadic updating circuit during grasp preparation.

Eye-hand coordination is relatively understood in terms of the transport component of reach, but little is known about the integration of saccade and visual signals for updating grasp configuration across eye movements. We set out to identify a putative human grasp updater and found a remarkably consistent cluster of regions including SMG and aIPS/SPL, (likely) receiving oculomotor inputs from prefrontal eye fields. This network provides the necessary neural machinery to integrate object features and saccade signals, and thus ensure grasp plans remain updated and coordinated with gaze-centered reach transport plans (2, 3). These new findings have several general implications: First, this circuit might explain some of the various symptoms of apraxia that results from damage to the posterior parietal cortex (61, 62). Second, the role of the inferior parietal cortex in both transsaccadic perception (20) and grasp updating supports the notion that inferior parietal cortex (a very late phylogenetic development) has high-level visuospatial functions for both ventral and dorsal stream vision (25). Finally, the various roles of specific parietal modules in spatial updating (63), visual feedback corrections (53), and (here) a combination of the two for action updating, support a general role for parietal cortex for detecting, differentiating, and compensating for internally and externally induced spatial perturbations.

## Methods

The York University Human Participants Review Subcommittee provided pre-approval for the study. Seventeen healthy, right-handed individuals aged 22-32 participated. However, due to excessive head motion, four of these participants were excluded. This left thirteen participants’ data for analysis, sufficient to achieve a power value of 0.987. All participants provided written consent and were remunerated financially for their time. Participants were placed supine within an apparatus equipped to track eye and hand motion (Figure 1*A*) while they performed a reach-to-grasp task (Figure 1*B*) in an MRI scanner. We used a 3 T Siemens Magnetom TIM Trio scanner to collect anatomical and functional data. BrainVoyager QX (Brain Innovations) was used to preprocess and analyze these data. For details, see *SI Materials and Methods*.

## Acknowledgments

We thank Joy Williams for her help in collecting the data as the MRI technologist, and Dr. Xiaogang Yan and Saihong Sun for technical and programming support. This work was supported by a Natural Sciences and Engineering Research Council (NSERC) Discovery Grant. During this study, B.R.B. was supported by the NSERC Brain-in-Action CREATE Program, and the Ontario Graduate Scholarship: Queen Elizabeth II Graduate Scholarships in Science and Technology. S.M. was supported by the Canadian Institutes of Health Research during experiments and later a Marie Curie Fellowship. J.D.C was supported by the Canada Research Chair Program.

