## Supplementary Materials and Methods for "Parietal Cortex Integrates Object Orientation and Saccade Signals to Update Grasp Plans"

**This PDF file includes:**

Supplementary text
References for SI reference citations

**Supplementary Information Text**

**Materials and Methods**

**Participants.**

To determine the appropriate number of participants required for a sufficient level of power in this study, we reviewed the most relevant previous literature and then did a power analysis. The current experimental design was based on our previous fMRI studies of transsaccadic memory (20) and grasp orientation (34). Thirteen participants were analyzed in the Dunkley et al. (2016) study and 14 in the Monaco et al. (2014) study. We have found previously that motor behavior increases cortical activation in the posterior parietal areas of interest for this study (1, 2), so we based our power analysis on the Monaco et al. (2011) study. We then chose the region of interest from that study that was closest to the posterior parietal activation we predicted. Specifically, we used the effect size (1.27) from their left posterior intraparietal sulcus (pIPS) activation, along with the following parameters: 1) two-tailed t-test option, 2) an $\alpha$ value of 0.05 (we had planned for one contrast), and 3) a high power value (0.98). Using these values in G*Power (Faul, Erdfelder, Lang, & Buchner, 2009), we calculated that 13 participants would provide a sufficient actual power value (0.987).

In order to obtain a reliable dataset of thirteen participants (after the exclusion criteria described in the analysis section below) we had to test seventeen participants. These were all graduate students from York University, Toronto, Ontario, Canada, with no known neurological disorders and normal or corrected-to-normal vision. These participants were all right-handed and were of 26.5 +/- 3.7 years of age (from 22 to 32). All participants provided written consent and were compensated financially for their time. The York University Human Participants Review Subcommittee approved all experiments.

**Experimental set-up and stimuli.**

Participants were asked to fill out an MRI screening form. Upon passing MRI screening, participants were informed about the task. Once they felt comfortable with what the experiment entailed, they were asked to assume a supine position on the MRI table, with their head in a six-channel coil tilted forward at a 20° angle (in order to allow for direct visibility of the objects) (51). To obtain a complete signal, we also placed a four-channel coil anteriorly on the head (51).

This experiment was conducted in complete darkness. In our set-up, we had red fixation light emitting diodes (LEDs) for participants to focus on during the entire duration of a trial. A fixation LED was placed to the left and right of the central stimulus (between 10-12° from the center of the stimulus to each LED; (51)). There was also a white LED that was used to illuminate the stimulus only when participants would grasp at a particular time point in each trial (Fig. 1*A*, *B*). These LEDs were mounted onto a rotatable platform that was placed above each participant's pelvis. LEDs were held in place by MRI-compatible rigid tubes (which were made of many units to allow for movement of the overall tube in order to position the LEDs accordingly).

The stimulus that participants had to grasp was a six-degree long bar with rounded ends (Fig. 1*A*, *B*) and centered on the platform. The bar could be rotated, but two MRI-compatible pins were placed in the surrounding area to ensure that the bar could only be oriented horizontally (0°) or obliquely (135°).

For each participant, right eye position was recorded using an infrared (IR) camera affixed to the right side of the MRI table (Fig. 1*A*). Eye movement signals were recorded using iViewX software (SensoMotoric Instruments) for offline analysis. We recorded, using a hand camera (Fig. 1*A*), the reaching and grasping movements of participants during each trial of every run.

Lastly, in order to reduce any motion artifacts in the imaging data, participants' upper arm and shoulder were immobilized using an MRI-compatible belt that was strapped down across their torso. Participants reached with their right hand and pivoted only from their elbow joint, with only the minimal rotation of the shoulder joint. Their right arm was supported with foam padding and sand bags to provide a comfortable height from which the arm could reach and grasp the object for the duration of the experiment. We also made sure that the addition of the padding was appropriate and allowed participants to reach and grasp the object appropriately.

**General paradigm/procedure.**

**Experiment.**

We used an event-related fMRI design to identify cortical areas involved in updating grasp orientation across saccade eye movements. To test this, we developed a behavioral paradigm that would provide interactions between transsaccadic orientation memory (20) and changes in the required grasp orientation (34). First, participants were placed in the MRI bore and a comfortable reach distance was determined for placement of the grasp stimulus by moving the platform along the MRI table. Participants were then trained to reach and grasp this bar stimulus (see previous) in response to a specific ‘go’ signal (Fig. 1*A*, *B*). At the beginning of each reach trial, participants rested their arm, bent at the elbow, on their abdomen in a position that was within comfortable reaching distance of the stimulus. Participants were instructed to use all digits of their right hand to grasp center of the object (Fig. 1*A*). Upon completing the grasp, participants returned their arm to the same resting position as prior to the reach.

Each trial started with the illumination of one of the two LEDs to the right and left of the central target. Then, the central target was illuminated for 250 ms. The target could be oriented at 0° or 135° (pseudorandomized and counterbalanced within and across runs). Participants were required to keep fixating for another 1.75 s. This first two-second phase was referred to as the ‘*Stimulus Presentation*’ phase (Fig. 1*B*). After this period, participants kept fixating on the same LED for another 1.75 s (‘Fixate’ condition) or make a saccade to the other LED, which would be illuminated while the previous LED would be extinguished (‘Saccade’ condition). Following this 1.75 s period, the object was illuminated for 250 ms. This was referred to as the ‘*Action Preparation*’ phase (Fig. 1*B*), when participants were expected to retain stimulus location and orientation information, and use this to prepare for a movement (1-2, 34). The object could now be oriented in the same orientation as in the first illumination/presentation (‘Same orientation’ condition, e.g., 0° orientation first and then, another 0° orientation; same for the 135° orientation) or a different orientation as compared to the first (‘Different orientation’ condition, e.g., 0° orientation first, followed by a 135° orientation and vice versa). Participants were then given 4 s to reach out to grasp the object in its final orientation as described above (‘*Action Execution’* phase) while still fixating the illuminated LED. Following this phase, the LED was set up for the next trial and participants had 16 s to rest while maintaining fixation (intertrial interval, ITI), so as to allow the BOLD signal to come back to baseline. The illumination of the stimulus marked the beginning of each trial and the end of the 16 s period of relaxation marked the end of the trial.

In order to create the different orientation conditions, one experimenter rotated the stimulus as needed in the scanner room, but out of the participant's view and in complete darkness. To reduce the possibility of participants predicting Different versus Same orientation based on sound feedback, the experimenter moved the stimulus away and back to its required orientation (also during Same orientation conditions).

The design of the experiment consisted of a 2 (Gaze Position: Fixate or Saccade) x 2 (Gaze Fixation Location: Left or Right) x 2 (Object Orientation: 0° or 135°) design. This produced eight condition types, which were repeated four times within one run. There were six runs in total. As mentioned previously, the condition types were pseudorandomized and intermingled within each run and across runs.

Compared to our previous study (20), we used a shorter stimulus period (total of 2 s for each stimulus presentation) in order to match an acquisition time of 2 s, and to ensure a reasonably long run/ experiment (given that a long ITI is needed to allow the BOLD to return to baseline). Recent studies have suggested that this transsaccadic integration can occur on the order of tens of ms (3, 16, 18). In addition, we chose a fixed ITI (no jitter) because we did not investigate response timing in this study and we wished to maximize our statistical power in order to detect transsaccadic integration signals (20).

**Saccade Localizer.**

To determine which regions are involved in the production of saccadic eye movements, we used a localizer that had a sequence similar to that of the experimental runs. This localizer comprised alternating periods of fixation and saccadic eye movements. First, a baseline of activity would be established as a result of participants fixating the illuminated LED for 18 s (two runs total of data were collected, where participants fixated the left LED first and right LED second, or vice versa). Then, every second for 6 s, the LEDs would alternate in illumination, resulting in saccades. After this, participants would then fixate the initial LED for 16 s. This fixation-saccade sequence was repeated eight times in the localizer run. There was a last fixation period of 18 s. For each of these two runs, 98 volumes were acquired, with 35 slices per volume (TR= 2000 ms, TE= 30 ms; in-slice resolution of 3 mm x 3 mm; slice thickness= 3.5 mm, no gap).

**Imaging Parameters.**

We used a 3T Siemens Magnetom TIM Trio magnetic resonance imaging (MRI) scanner. The functional experimental data were acquired using an echo-planar imaging (EPI) sequence (repetition time [TR]= 2000 ms; echo time [TE]= 30 ms; flip angle [FA]= 90 degrees; field of view [FOV]= 192 x 192 mm, matrix size= 64 x 64 with an in-slice resolution of 3 mm x 3 mm; slice thickness= 3.5 mm, no gap) for all six functional runs in an ascending and interleaved manner. Along with functional data, a T1-weighted anatomical reference volume was acquired using an MPRAGE sequence (TR= 1900 ms, FA= 256 mm x 256 mm; voxel size= 1 x 1 x 1 $mm^3$). For each volume of anatomical data obtained, 192 slices were acquired. We collected 395 volumes of functional data for the experimental runs. Each volume comprised 35 slices.

**Analysis.**

**Behavioral data.**

We monitored eye position during the experiment and analyzed it offline, to verify that participants fixated on the appropriate LED and did not make any additional, unnecessary saccades during trials. Any trials showing inappropriate fixation or saccades were removed from additional analysis. Similarly, video data were analyzed offline to determine if the participant grasped the object at the required time. Any trials during which any anomaly in grasping occurred (i.e., participant grasped the object too early or too late, etc.) were removed from further analysis by being designated as confound predictors in the general linear model (see below). Overall, eight trials were removed from the entire data set (two trials each were excluded from two participants and one trial from another four participants).

**Functional imaging data: Experimental.**

A general linear model (GLM) was created for each run for each participant. A predictor was used as a Baseline for the period of fixation at the beginning and the end of each run ("Baseline"), accounting for the first 18 s of each run and the 16 s inter-trial interval. The initial 2 s *(Stimulus Presentation* phase) during which the object was illuminated and participants had to fixate an LED was assigned a predictor that indicated the location of the fixation LED ("Adapt_LVF" and "Adapt_RVF" if the fixation was on the right LED or left LED, respectively; left and right visual field for LVF and RVF, respectively). The subsequent *Action Preparation* phase (2 s) was assigned one of four predictors: “Sacc_NovFeature”, “Sacc_RepFeature”, “Fix_NovFeature”, or “Fix_RepFeature” for when participants made a saccade or fixated and, for each of these, whether the orientation of the object was the same (Same orientation condition) or different (Different orientation condition). The *Action Execution* phase was divided in two 2-s phases. There were four predictors for the first 2 s of the grasp event. These predictors were based on the direction of the preceding saccade and upon whether the orientation of the object that was being grasped was the same in the *Action Preparation* phase as in the *Stimulus Presentation* phase (Same orientation condition) or different (Different orientation condition). Thus, the four predictors were: “Motor Execution_Sacc_NovFeature”, “Motor Execution_Sacc_RepFeature”, “Motor Execution_Fix_NovFeature”, and “Motor Execution_Fix_RepFeature”. The following 2 s of the *Action Execution* phase were provided a “Motor Execution” predictor. These predictors comprised each GLM for each participant (BrainVoyager QX 2.8, Brain Innovation). Each predictor variable was convolved with a haemodynamic response function (standard two-gamma function model) (26). GLMs were modified through the addition of confound predictors for eye movement or hand movement errors. If a GLM had more than 50% of the trials being modelled in the confound predictor, the GLM for that run was not included in the overall population level GLM (random effects GLM, RFX GLM).

Additionally, functional data for all runs across all participants was preprocessed (slice time correction: cubic spline, temporal filtering: <2 cycles/run, and 3D motion correction: trilinear/sinc). Data for runs that had abrupt motion of over 2 mm were excluded from the RFX GLM and additional analysis. As a result, four participants' data were excluded because more than half of the runs were unusable due to abrupt, excessive head motion of over 2 mm. The anatomical data was transformed to a Talairach template (4) and the functional data from the remaining 13 participants were coregistered using gradient-based affine alignment (translation, rotation, scale affine transformation) to raw anatomical data. Lastly, functional data were smoothed using an FWHM of 8 mm.

We used voxelwise analysis of β-weights obtained during the *Action Preparation* and *Action Execution* phases in order to determine whether there are any cortical regions that are modulated by saccades during the period of preparation and by transsaccadic integration of object orientation during grasping. β-weights were extracted from the selected regions and were used to test our hypotheses. For each *a priori*-motivated t-test conducted within hypothesized regions, we indicate the relevant statistical values (i.e., t-statistic and p-values).

**Functional Imaging Data: Localizer.**

For the preprocessing of functional data for the localizer, see above section. On the basis of excessive head motion (>2 mm), one person's data was completely excluded, half of the data was excluded for a second person (runs where initial fixation was on the left), and half was excluded for a third person (runs where initial fixation was on the right). Using the remaining functional data, we ran an RFX GLM on the data for each of the localizers. For the saccade localizers, we had three predictors: a 9 s

“Baseline” predictor, a 16 s fixation “Fix” predictor, and a 6 s saccade “Sacc” predictor. The results of the saccade localizer were used to identify which regions are involved in saccade production in our task specifically (Figs. 2*B*, 4*B*).

**Functional Connectivity: Psychophysiological Interaction Analysis.**

In order to determine the network of cortical regions that interact to update saccade signals during the preparation of a grasp, we conducted psychophysiological (PPI) analysis (5-7) during the *Action Preparation* phase. In order to carry out this analysis, we had three predictors: 1) physiological component (z-normalized time courses obtained from the seed regions for each participant for each included run), 2) psychological component (predictors of the model were convolved with a haemodynamic response function), and 3) psychophysiological interaction component (multiplication of z-normalized time courses with task model in a volume-by-volume manner). For the task model produced for the psychological component, the Saccade predictors were set to a value of ‘+1’, whereas the Fixation predictors were set to a value of ‘-1;’ all other and baseline predictors were set to a value of ‘0’. Single design matrices (SDMs) were created for each participant for each included run. These were subsequently included in an RFX GLM for right SMG (5) in order to determine functional connectivity with each of these regions.
